# Cellular dynamics in the maize leaf growth zone during recovery from chilling depends on the leaf developmental stage

**DOI:** 10.1101/2023.08.17.553746

**Authors:** Cindy M.S. Lainé, Hamada AbdElgawad, Gerrit T.S. Beemster

## Abstract

*Zea mays*, a major crop, is highly sensitive to chilling which frequently occurs during its seedling stage and negatively affects yields. Although the direct effect of chilling is well-studied, the mechanisms determining the subsequent recovery are still unknown. Our goal is to determine the cellular basis of the dynamic leaf growth response to chilling and during recovery of leaves exposed before or after their emergence. We first studied the effect of a 3-day cold spell on leaf growth at the plant level. Then, we performed a kinematic analysis to analyse the dynamics of cell division and elongation during recovery of the 4^th^ leaf after exposure to cold before or after emergence. Our results demonstrate that cold more strongly reduced the final length of non-emerged than emerged leaves (-13 vs -18%). This was not related to growth differences during cold, but a faster and more complete recovery of the growth of emerged leaves. Kinematic analysis showed that this difference was due to a higher cell division rate on the 1^st^ and a higher cell elongation rate on the 2^nd^-day of recovery, respectively. The dynamics of cell division and expansion during recovery determine developmental stage-specific differences in cold tolerance of maize leaves.

## Introduction

### Cold spells reduce maize production

Despite future climate scenarios predicting higher average temperatures, cold spells, and short periods of successive cold days, are a major threat to early-sown crops in Europe. Such cold spells not only affect the plant at the seedling stage but result in a lower yield in crops, including maize (*Zea mays*; Frei, 2000), cotton (*Gossypium hirsutum* L.; Pettigrew, 2002) and soybean (*Glycine max* L; Toda et al., 2011). As a sub-tropical species originating from Mexico, Maize (*Zea mays*) is particularly sensitive to cold (Greaves, 1996). Nevertheless, farmers can benefit from early sowing in temperate climates by extending the length of the growing season and thereby increasing yield or by facilitating early harvesting to allow a second crop to be sown in the same growing season. Early spring sowing, however, increases the risk of exposure to transient cold spells.

### Effect of temperature decrease

Temperature is an important factor affecting physiological processes, including photosynthesis and growth in maize (Ben-Haj-Salah and Tardieu, 1995; Giauffret et al., 1995; Cholakova and Vassilev, 2017). It is the primary determinant of leaf emergence and leaf elongation rate (Granier and Tardieu, 1998). Consequently, the accumulated thermal time (degree days) is often used to analyse maize growth and development instead of chronological time (days) (Wang, 1960; Granier and Tardieu, 1998). As temperature decreases (down to a temperature base of 8°C for maize), most physiological processes slow-down (Birch et al., 1998). Below this threshold, plant growth stops and cold stress occurs, damaging the photosynthetic system and inducing oxidative stress (Greaves, 1996; Farooq et al., 2009). However, our recent meta-analysis demonstrated that 8 degrees is not an absolute boundary and growth still occurs below this threshold (Lainé et al., 2022). The effect of cold, so-called chilling stress, defined as exposure to suboptimal cold temperatures above 0 °C, has been well-studied in many crops and especially maize seedlings (Lainé et al., 2022).

Maize plants are capable of efficient recovery even from severe stress conditions such as flooding (Yeung et al., 2018), drought (Vilonen et al., 2022) or heavy metal (Chmielowska-Bąk and Deckert, 2021). However, only a few studies (e.g. Avila et al., 2018) investigated the recovery after chilling in maize and we are unaware of studies investigating the dynamics of leaf growth recovery. In cotton, the relative growth rate (RGR) and Net Assimilation Rate (NAR) of seedlings exposed to 1-day of cold fully recovered after 3-days, but this was not the case after exposure to 3-days of cold (DeRidder and Crafts-Brandner, 2008). Similarly, Atkinson et al (2015) observed that RGR and NAR of *A. thaliana* seedlings decrease by 70% during exposure to 5°C, while during recovery RGR increases by 50% due to a partial recovery of the NAR (68% of the control) and nitrogen productivity (37% of the control) and full recovery of the plant nitrogen content (PNC). This suggested that growth recovery after cold is likely due to NAR. Surprisingly, NAR of 14-day cold-treated tomato seedlings at 6°C did not recover after 7 days in control conditions (Brüggemann et al., 1992). In contrast, after 2-days recovery, stomatal conductance substantially increases. Leaf area and shoot fresh weight (FW) of 28-day cold-treated tomato seedlings slowly recover during 14 days and substantially increases from 14 to 21 days recovery (Brüggemann et al., 1992). In maize, increasing Rubisco levels in mature maize leaves improved growth by maintaining a higher photosynthetic rate during exposure to cold, which helps to recover faster from cold and reduces damage in the photosynthetic system II (Salesse-Smith et al., 2020). This suggests a dependence on restored carbohydrate supply from the mature leaves to the growing tissues. Interestingly, Garbero et al (2012) show that, in the chilling sensitive pangola-grass, a higher cold tolerance level was associated with a quicker recovery of leaf dry weight and FW, which was likely due to the observed increase in auxin (IAA) during the period of cold stress. These observations suggest that restoration of a high growth rate at which plant recovers, and the short period taken characterise a better recovery.

### Developmental stage specificity of cold tolerance

At the plant level, cold affects growth differently when occurring at contrasting developmental stages. At 6°C, maize is able to germinate (Miedema, 1982). In contrast, the areal part of the plant is especially sensitive to the same temperature during the seedling/vegetative stage when plants pass from the heterotrophic to the autotrophic stage (Greaves, 1996). Leaf growth could also be affected differently depending on the timing of exposure to cold at different leaf stages (De Vos et al., 2020). Important physiological changes exist between an emerged leaf and a leaf that is surrounded by the whorl of older leaves. Before emergence, primordial growth is exponential (Beemster and Masle, 1996) and most of its resources are allocated to growth. Once the leaf has emerged, it is exposed to light and actively starts photosynthesis. At present, it is unclear how the developmental stage of maize leaves affects their response to cold and subsequent recovery. Responses to abiotic stress can differ when occurring at different leaf stages. For instance, in *Arabidopsis* young leaf tissues have a higher plasticity in traits such as growth responses, a more efficient NPQ activation and DNA repair (Rankenberg et al., 2021). The loss of this plasticity with age corresponds with a decline in stress resilience of older tissues but can be compensated by the acquisition of physical (e.g., cuticle) or chemical (e.g., anthocyanins) defences (Rankenberg et al., 2021). Therefore, we expect that responses to chilling stress at different leaf developmental stages, particularly before and after emergence will differ.

### Kinematic analysis of cell division and expansion during leaf growth

Leaf growth is driven by two processes: cell proliferation (where cells are produced by cell expansion in combination with mitosis) and cell elongation (where they expand to their mature size in the absence of mitosis; Green, 1976). It is frequently assumed that during the linear phase of leaf growth after its emergence, the growth is in a steady-state where leaf elongation rate is constant, cell length profile and length of the growing zones are stable, as observed in maize (Rymen et al., 2007; Aydinoglu, 2020) and sorghum (Bernstein et al., 1993). Assuming a steady-state of growth simplifies kinematic analysis, a well-developed method used to determine the contribution of cell division and cell elongation to organ growth (Bernstein et al., 1993; Beemster and Baskin, 1998; Fiorani and Beemster, 2006). When cold stress occurs, those two processes are inhibited, resulting in the slowing down of the growth of the leaf as a whole. Cold nights (25°C day/4°C night as compared to 25/18°C for control, reduces meristem activity, while cell elongation appears to be less affected in maize (Rymen et al., 2007; Aydinoglu, 2020). To date, no kinematic analysis has been performed on leaves with cold stress applied during both night and photoperiod. The impact of cold stress occurring during the photoperiod is probably more severe than during the night as it also perpetuates chloroplast development, leading eventually to cell death (Allen and Ort, 2001; Gómez et al., 2004).

Therefore, our research aims to understand the cellular basis of leaf growth responses of maize seedlings to cold during different stages of development, i.e., before and after leaf emergence, and during recovery under control conditions. Our first objective is to define the impact of a transient period of 3 days of cold on leaf growth rates during different stages of leaf development. Our second objective is to understand the contribution of cell division and elongation in leaves of different developmental stages in response to cold and during subsequent recovery using kinematic analysis.

We tested whether (i) cold differently affects the growth of leaves in different phases of their development (emerged, about to emerge, initiated; Atkinson et al., 2015). As it was observed that the direct effect of cold stress mainly reduces cell division rate (Rymen et al., 2007) we hypothesise (ii) that the direct effect of exposure to cold is similar between leaf developmental stages (D0 and D3 plants) and is mainly caused by a reduction in cell division rate. (iii) Unlike the direct effect of cold, we expect that both cell division and elongation contribute to cold recovery, which differentially affects the growth of leaves from different developmental stages.

## Materials and Methods

### Plant growth

We used the maize (*Zea mays*) inbred line B73 to study the effect of cold on leaf growth. Plants were grown in a growth chamber set up at 24/18°C with a 16/8h light cycle, 300–400μE m^−2^ s^−1^ photosynthetically active radiation, provided by high-pressure sodium lamps and 50% humidity. For cold treatment, maize plants were transferred from the growth chamber to a growth cabinet set up at 8/4°C for a 16/8h light cycle and 50% humidity for 3 days and transferred back to the growth chamber for recovery. The cold treatment has a considerable effect on plant growth and represents a realistic cold spell in Western European spring. Control plants remained in the growth room as preliminary experiments showed there was no growth cabinet effect on Leaf Elongation Rate (LER) under control conditions. To study the effect of cold on different developmental stages of leaf number 4, we applied cold treatments just before leaf emergence (D0 plants) and 3 days after (D3 plants) (**Fig. 1**). We harvested 6-7 leaves per treatment for kinematic analysis in the middle of the photoperiod. A leaf that had just emerged was considered a D1 leaf once its appearance could be observed above the whorl of older leaf sheaths. A subset of 13-15 plants per treatment was used to analyse leaf emergence and measure the length of the 4^th^ leaf until it reached its final size.

**Figure 1:**
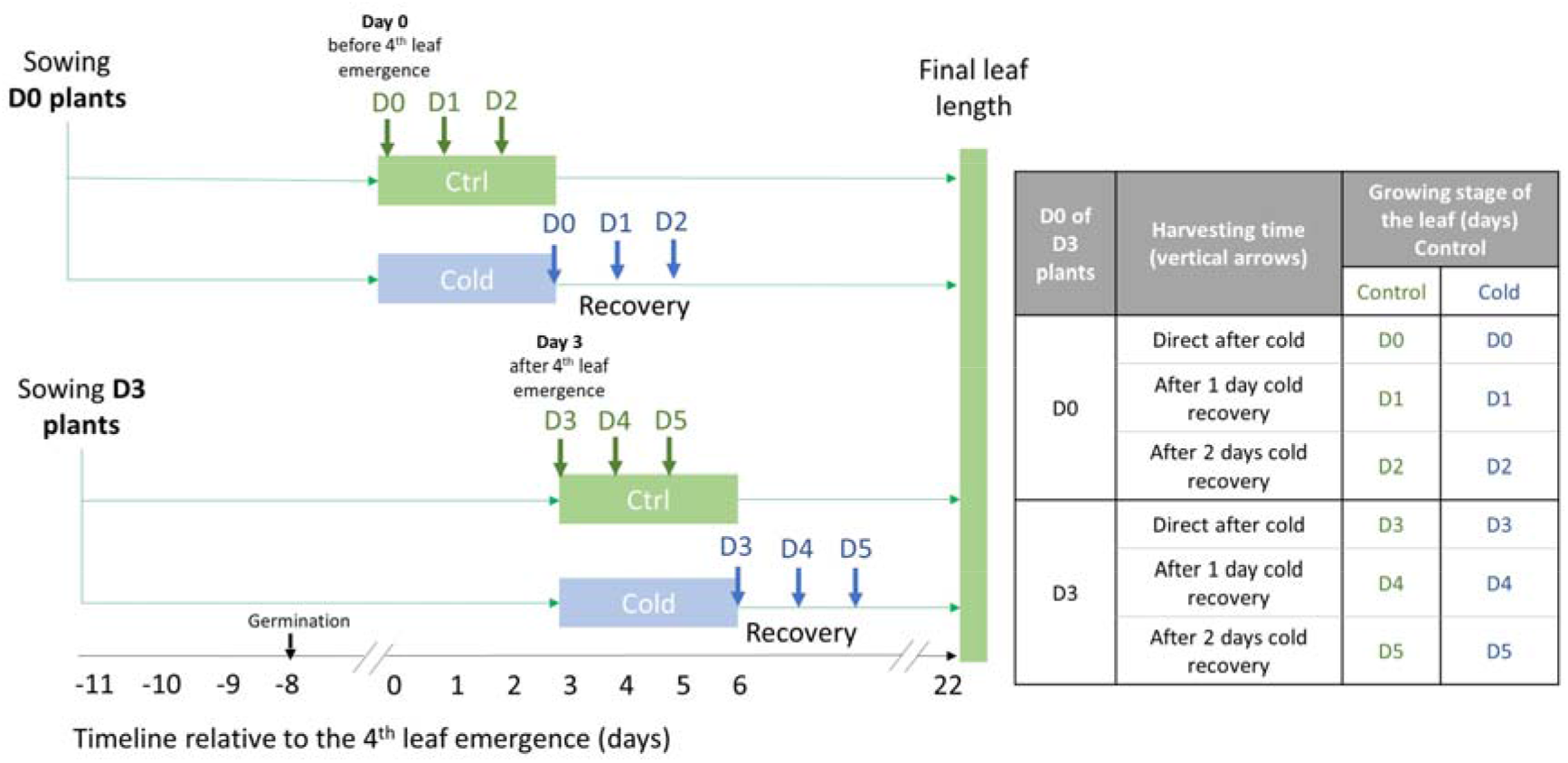
Overview of the experimental setup to study the impact of a transient cold before and after leaf emergence on maize leaf growth. Timing relative to leaf emergence of sowing, temperature treatments and harvests for kinematic analysis (arrows). By harvesting directly after 3 days of cold, as well as 1 and 2 days after cold, we aim at studying the direct effect of cold and subsequent recovery. To compare cold-treated leaves with controls at the same growth stage, we harvested the control plants at the time cold treatment started. Assuming that during cold development essentially stops, developmental stage (Dn) is therefore delayed by the duration of the cold treatment. The cold-treated **D0 plants** were transferred to cold just prior to 4^th^ leaf emergence, while the **D3 plants** were placed in the cold 3 days after 4^th^ leaf emergence. For kinematic analysis, 6-7 plants were harvested per treatment. For the final leaf length measurement, 13-15 plants were measured.

### Phyllochron rate and phyllochron index

The concept of phyllochron, based on the emergence rate of new leaves in maize, is nearly constant during the seedling stage, when expressed in thermal time units (°Cd, degree-days) (Birch et al., 1998). Maize thermal time (growing degree days, GDD) is calculated with the following formula = ((Daily Max Temp (day) x 16h + Daily Min Temp (night) x 8h) / 24) – Tbase (8°C). However, as we know that maize plants continue growing even below 8°C (Lainé et al., 2023), we decided to also include the growth parameter relative to the time in days for our research in the supplemental data (**Fig. S2**).

The phyllochron rate (°C Day. leaf^-1^) is the rate of leaf emergence. The phyllochron corresponds to time intervals between the emergence of two successive visual leaves (Birch et al., 1998; Plancade et al., 2022). It is obtained from the relationship between the number of leaves and the growing degree per day. The phyllochron index (PI) is a measure of the plant age interpolating the period between emergence of successive leaves:

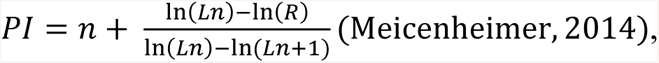

where n is the sequential index number of leaves for which the PI is being calculated, with n increasing in an acropetal direction, R is the reference leaf length, Ln is the length of a leaf that is equal to, or longer than R, and Ln+1 is the length of a leaf that is shorter than R. In **Fig. S1a,** the reference value of leaf length (R), i.e., 35cm, is designated with a dashed horizontal line.

The reference length (R) must be such that the length of leaf n is equal to or greater than the reference length, while the length of leaf n+1 is less than the reference length (Ade-Ademilua et al., 2005). For this reason, we choose a reference length of 35cm. All leaves developed before leaf n must be longer than the reference length (Ade-Ademilua et al., 2005).

### Leaf Elongation Rate

To obtain the Leaf Elongation Rate (LER) of leaf 4, we daily measured its length (LL) relative to the pot surface with a ruler between 9 and 10 AM. LER is calculated by taking the change in leaf length between two successive days, using the following formula: (LL _d1_ - LL_d0_)/24 and is expressed in mm per hour. Leaf 4 of D0 plants could not be measured as the leaf wasn’t visible yet. Thus, a set of 15 plants per treatment were grown and harvested at D-1, D0, and D1 relative to leaf emergence. Leaf 4 was dissected, and its average leaf length was used to calculate LER for D0 and D1 plants. This LER did match the LER when using the length relative to the pot surface.

### Meristem and cell length profile

Meristem size and cell length measurement were measured according to the protocol of Sprangers et al 2016.

### Data and statistical analysis

To determine the effect of cold exposure on D0 and D3 on the phyllochron we tested whether the slopes of linear regression of PI were significantly different from each other using ANCOVA.

The kinematic analysis of leaves uses three datasets (leaf elongation rate, meristem length, and cell length profile) to calculate different growth parameters at a cellular level. Data were analyzed in R software (http://www.Rproject.org/) using the leafkin package (Bertels and Beemster, 2020). A more detailed explanation of the kinematic analysis can be found in the protocol developed by (Sprangers et al., 2016).

Under non steady-state conditions, density changes in the division zone need to be taken into account to calculate the cell division rate (D’’) and cell production rate (P’’) (Silk, 1992; Fiorani and Beemster, 2006). To determine, rate of density change (ΔDensity) in a constant meristem we used calculated the derivative of a 3 or 5 points polynomial fitted to the data of cold treated and control plants, respectively:

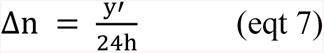

Where y’ (y’=2ax+b) is the derivative of the polynomial equation (y=ax^2^+ bx +c).

Once the delta number of extra cells produced each day was obtained, we used it to re-calculate the cell production rate (P’’) and cell division rate (D’’) under non steady-state conditions (Erickson, 1976; Beemster and Baskin, 1998).

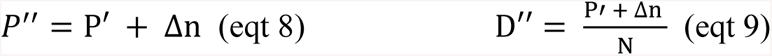

Where P’ is the cell production rate between 2 successive days, Δ_n_ is the number of cells produced in the meristem per hour and D” is the cell division rate at a non steady-state. This method was tested by comparing its output with the classical method during steady growth. More details information can be found in **Method S1.**

We performed 3-way ANOVA, 2-way ANOVA, and Student’s T test using SPSS (IBM SPSS Statistics for Windows, Version 28.0) depending on the parameter from kinematic analysis and final leaf length measurement.

## Results

### Cold stress effect at the whole shoot level

We first determined the effect of a transient 3-day cold period on leaf growth of a maize seedling. To this end, we measured the length of all emerging leaves for 30 days after germination for control and cold-treated plants (**Fig. S1**). To determine the cold stress-specific responses of LER, time (days) was expressed in temperature-compensated units or degree days (°C. Day^-1^). As expected, cold had no effect on leaf growth dynamics (**Fig. 2a**) and final leaf lengths were similar to those of the controls (**Fig. 2b**) of leaves 1 and 2 that had completed their growth before the cold occurred, had.

**Figure 2:**
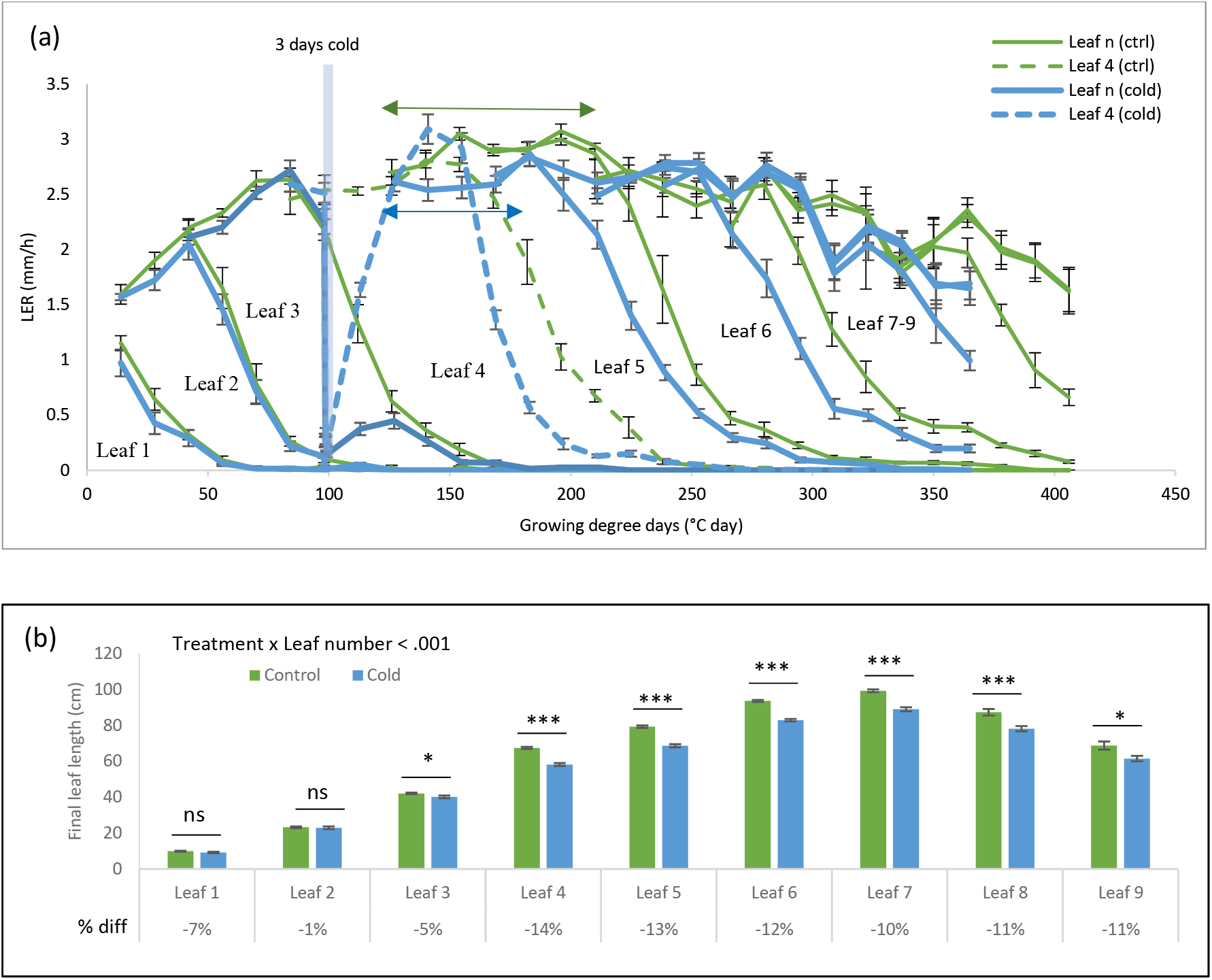
Effect of a transient cold on elongation rate and final length of maize leaves. Impact of a 3-day cold spell on leaf elongation rate (A) and final length (B) of the first 9 leaves at 30 days after germination. Maize plants were either grown in control condition (green) or exposed to 3 days cold on the third day after emergence of the 4^th^ leaf (blue). The blue band, in A, indicates the period of cold treatment. The arrow represents the length of the steady growth state of leaf number 5 for control and cold-treated plants. The x-axis for LER (a) represents the growing degree day expressed in thermal time units (°Cd, degree-days) (Birch et al., 1998). Data are shown as Mean ± SE with a sample size of n= 9 (p-value is indicated as ns: p >0.05; *: p < 0.05; **: p <0.01; ***: p <0.001).

In contrast, the growth of leaf 3 which was almost completed at the time of the cold treatment was strongly inhibited, resulting in a slight (-4%) but significantly reduced final leaf length compared to control (p=0.049). Leaf 4 was exposed to cold 3 days after its leaf emergence during steady-state growth. LER was reduced by more than 90% during the cold, recovering completely after. However, the growth phase was shortened by about 44-degree days, leading to a reduction of 12% in final leaf length (p< 0.001). Exposure to cold also affected the growth of leaf 5, the first to emerge after the cold. The duration of the growth phase was reduced (Arrows **Fig. 2a**), but also LER during steady-state growth was lower (**Fig. 2a**), resulting in a 13% of final length (**Fig. 2b**). Leaf 6 was similarly affected, while leaves 7 to 9 did not finish their growth during the experiment, remaining shorter at 30 days after germination, primarily due to the nearly complete developmental arrest during the cold.

### Effect of cold on phyllochron index

As we noticed a shift in the timing of the growth of individual leaves (**Fig. 2a**), we next investigated whether cold stress experienced during early growth of the maize seedling affects leaf emergence rate. To address this question, we determined the phyllochron index, which expresses the formation of new leaves at the shoot apex. Consistent with the early termination of leaf growth, the evolution of the phyllochron index is faster in plants after exposure to cold (**Fig. 3**).

**Figure 3:**
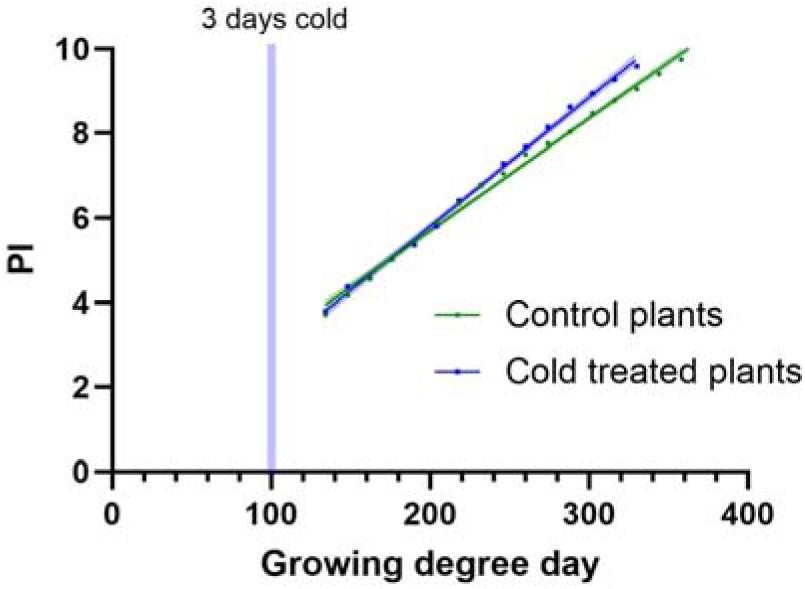
Impact of a transient cold on the phyllochron index of seedling *Zea mays* plants. Phyllochron index per growing degree day. Plants were either grown in control conditions at 24/18°C for 16/8h light/dark (green) or cold at 8/4°C with the same photoperiod (blue). Cold-treated plants were placed 3 days in the cold 3 days after 4^th^ leaf emergence (blue vertical band). The P-value from the ANCOVA test is p<0.001. Linear equation, p-value and R squared from T-test are for the control plants: y_ctrl_=0.027x+0.382, p<0.001, R^2^= 0.97 and the cold treated plants: y_cold_=0.030*x– 0.278, p<0.001 and R^2^= 0.97.

Those observations are consistent with the reduced duration of the leaf growth after cold stress (**Fig. 2a).** A correlation between faster leaf appearance and shorter mature leaves was also observed by Riva-Roveda et al. (2016) in maize and by Pablo et al. (2022) and in rice.

### Cold stress effects at the leaf level

After observing that a transient 3-day cold differently affects the growth of leaves in different positions, which are in different stages of their development, we decided to study how cold exposure during contrasting developmental stages of the same leaf (number 4) affects its growth. To this end, plants were placed in the cold either just before (D0) or at 3 days after 4^th^ leaf emergence (D3) and the growth of the 4^th^ leaf was compared with that of control plants of the same age.

Leaf elongation rate (LER) was reduced by 98% for D3 plants and 78% for D0 plants during cold (**Table S1**). Interestingly, although LER of D0 leaves was less inhibited by cold, they recovered more slowly than D3 leaves (**Fig. 4a**). D3 plants take only 3 days to recover 95% of LER, while D0 reached a maximum of 83% LER of controls after 6 days of recovery (**Fig. 4a**). As a result, mature leaf length was reduced by 13% and 18% for D3 and D0 cold-treated plants respectively compared to the control plants (**Fig. 4b**; p interaction treatment x timing = 0.017).

**Figure 4:**
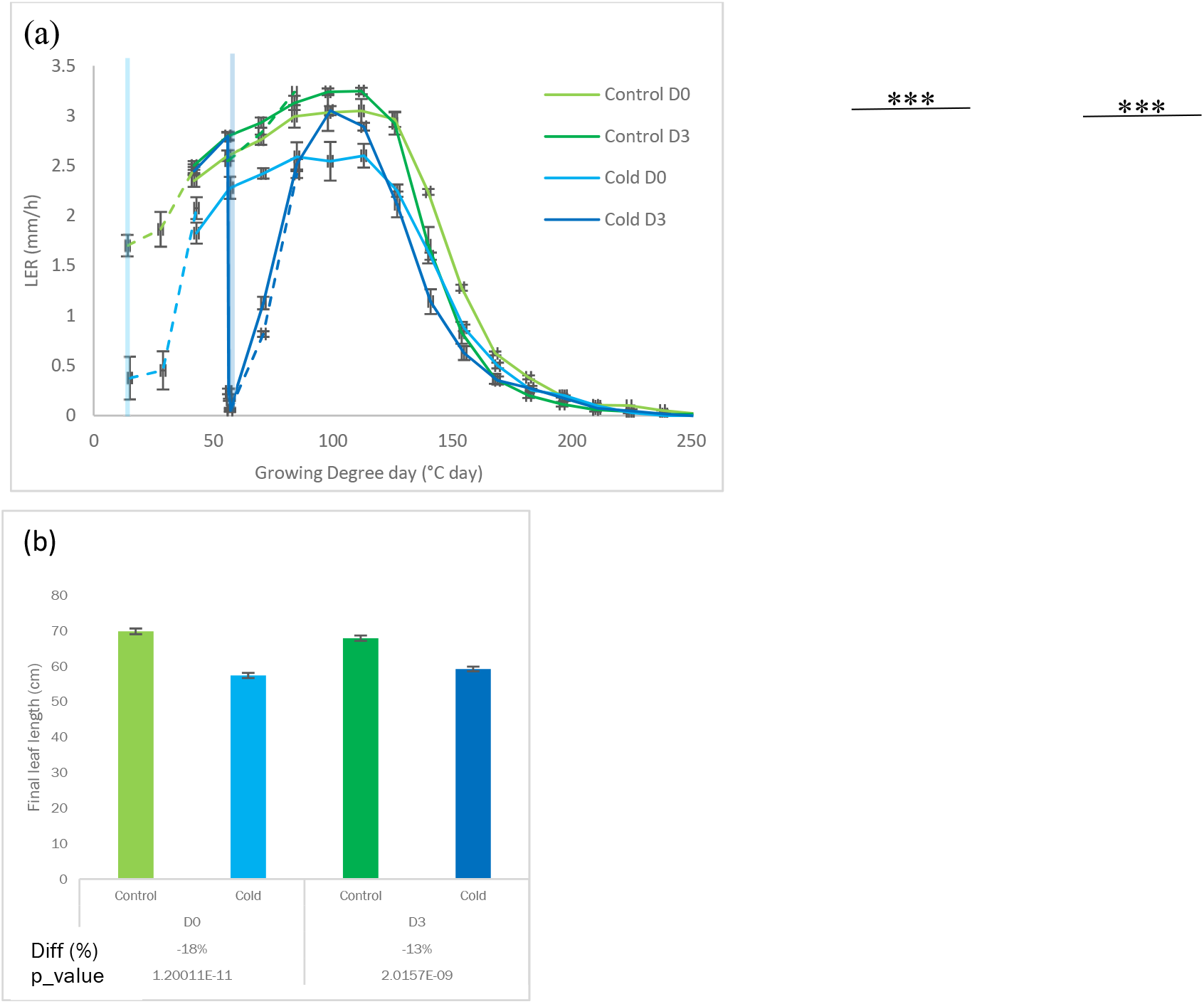
The effect of a 3-day cold period at different developmental stages on maize leaf growth. Time-course of leaf elongation rate from the 4^th^ leaf of maize in control (green) or cold condition (blue). Cold-treated plants were placed for 3 days in cold at D0 before 4^th^ leaf emergence (light) or at D3 after 4^th^ emergence (dark) and transferred back to control condition for recovery. (a) leaf elongation rate calculated from daily leaf length measurements in function of thermal time with 0 GDD being the first day of leaf measurement. (b) final leaf length. Interaction treatment x timing: p-value=0.017. Data are represented as mean ± SE, N=13-15. The dotted lines correspond to additional measurements on a separate experiment used for kinematic analysis.

### Kinematic analysis

At the cellular level, leaf growth (LER) is determined by cell division and elongation. To understand the cellular basis of different responses of D0 and 3 plants, we performed a kinematic analysis at three-time points: directly after 3 days of cold and after 1 and 2 days of recovery. The kinematic approach calculates cell division and expansion parameters based on measurements of LER, meristem size, and cell length profile (Sprangers et al., 2016; Bertels and Beemster, 2020). Commonly, kinematics is based on the assumption of steady-state growth, i.e. a stable leaf elongation rate, cell length profile and meristem size for several days (Muller et al., 2001; Fiorani and Beemster, 2006).

Because our experimental setup spans a 6-day time window to assess the effect of cold at D0 and D3, the controls represent a time series of the first 6 days after emergence. In contrast to the steady-state assumption, we observe that even under control conditions LER is not steady-state, but gradually increases during the first 6 days after emergence (**Fig. 4a; p = 0.001**).

### Cell length profile

To perform kinematic analysis, we first determined the cell length profile at the base of the leaf. For control plants, the length of the cells throughout the growth zone decreased over time (**Fig. 5**). This shows that, progressively, cells in the division zone divide at a smaller size, and this size difference is maintained in the elongation and mature zone.

**Figure 5:**
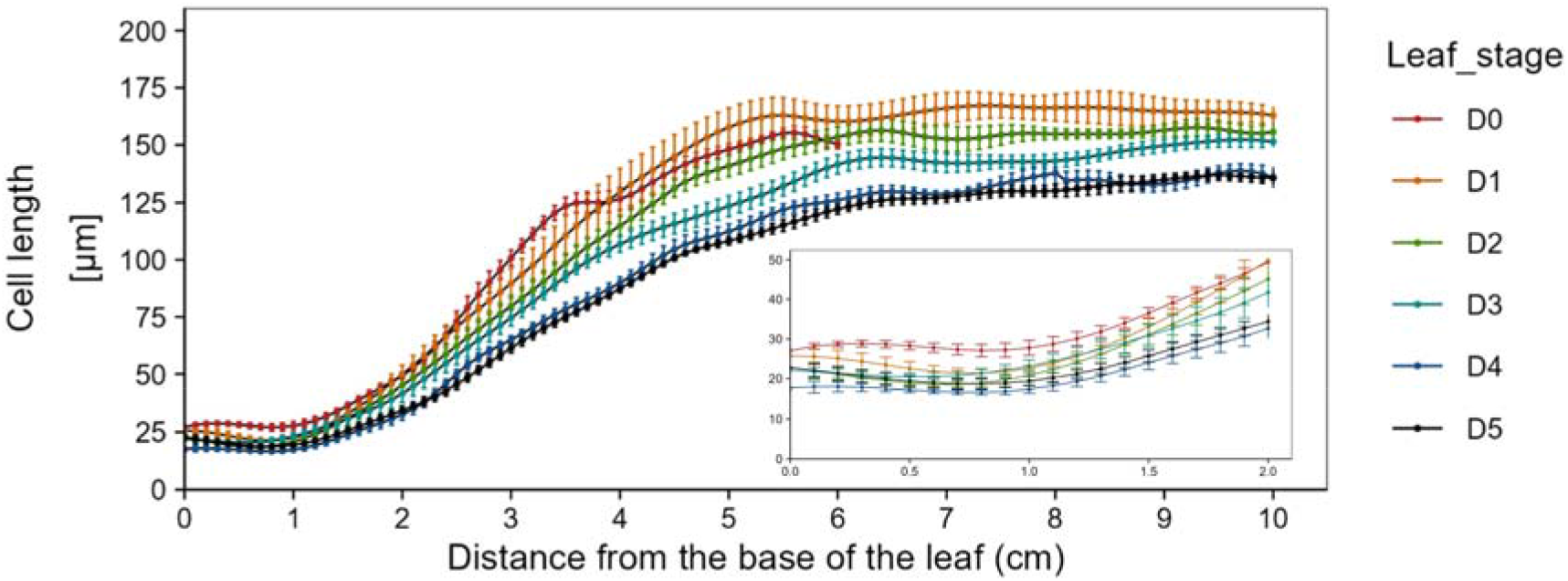
Evolution of the cell length profile of epidermal cells in the growth zone of the 4^th^ leaf of maize plants during the first 6 days after emergence under control conditions. The growth stages of the leaf range from the day before (D0) to 5 days after emergence (D5). Sample size: N=6-7, Error bars on the y-axis are confidence intervals at 95% around the mean length.

After cold exposure, the cell length profile of D0 and D3 leaves showed a similar response: The length of mature cells (l_mat_) was not affected, but similar to (Rymen et al 2007) cell size in the meristem was conspicuously reduced (**Fig. 6a, d, Table S1**). This suggests that expansion of proliferating cells is more inhibited by cold than division (Green, 1976). After 1 day of recovery, the difference in meristem cell size disappeared, suggesting that the expansion of proliferating cells recovers fast. At the same time, the cell length profile in the elongation zone of cold-treated plants was strongly affected by prior cold exposure. Cell size in the elongation zone of D0 and D3 treated plants was significantly smaller than in control plants from 2.5 cm onwards. Cell length of cold-treated plants continued to increase up to 7 and 8 cm for D0 and D3 treated plants respectively, while control plants reached mature cell length at 5 and 6 cm from the leaf base. Nevertheless, mature cell length (l_mat_) appeared unaffected (**Fig. 6b, e, Table S2**). After 2 days of recovery, the cell length profile became more similar to the controls, but l_mat_ was reduced by 20% for D0 (p=0.005) and 27% for D3 (p=0.005) (**Fig. 6c, f, Table S3**).

**Figure 6:**
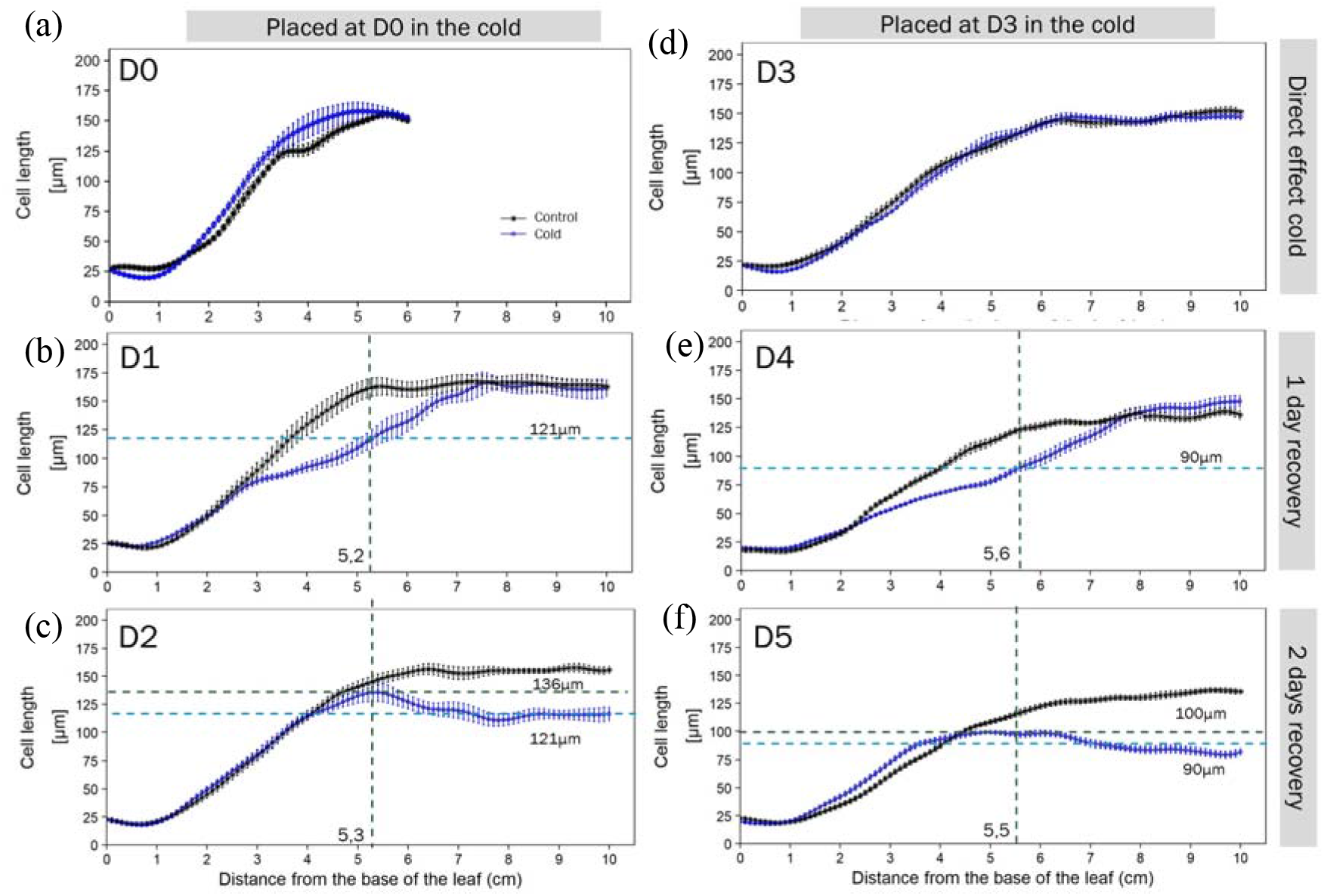
The effect of a 3-day cold period at different developmental stages on the cell length profile of the maize leaf. Plants were either grown in control (black lines) or cold conditions for 3 days (blue lines) at D0 (a-c) or D3 (d-f) after their emergence. Cold-treated plants were harvested directly after cold (a, d), after 1-day recovery (b, e), or 2 days recovery (c, f). The growth stage of the leaf is indicated with D0-D5 from 0 days before its emergence to 5 days after emergence. Sample size: N=6-7, Error bars on the y-axis are confidence intervals at 95% of the mean length.

These changes indicate that the cell length profile is not in steady-state. As a consequence, at D1 and D4, we are looking at a transient situation, where cells from 6 cm from the base onwards were already mature during the cold treatment. Upon resumption of growth, these cells are progressively displaced to more distal positions. The cells located in the elongation zone during the cold reach a mature cell size that is significantly lower than that of controls, forming the plateau on D2 and D5 (horizontal blue line, **Fig 6c,f**). Interestingly, l_mat_ at the boundary of the elongation and mature zone is higher than at larger distances, forming a consistent peak (horizontal black line **Fig 6c,f**), indicating an ongoing recovery of expansion in the elongation zone. Consistently, on D6 (**Fig. S3**), l_mat_ of D3 plants has recovered and is similar to the control ones. This implies that the lower plateau of mature cell size at 2 days of recovery corresponds to the size the elongating cells of the day before will reach (horizontal blue line) (**Fig. 6b, e**). Taking this into account, we define the length of elongation zone after 1 day of recovery as the position where cells reach the mature cell size observed in D2/5 (**Fig. 6b, e**). By doing so, the length of the growth zone of day-1 recovery of cold-treated plants is approximately 5.2 cm for D0 plants and 5.7cm for D3 plants.

### Spatial map of the leaf growth zone

Based on the localisation of (DAPI stained) mitotic cells, we determined the effect of developmental stage and cold on meristem size. Under control conditions, the meristem of leaves before emergence (D0) was substantially bigger than emerged leaves (22 vs 18 mm; **Fig. 7**). Once the leaf emerges (D1 to D5), meristem size remained approximately stable. In contrast, the elongation zone increased over time.

**Figure 7:**
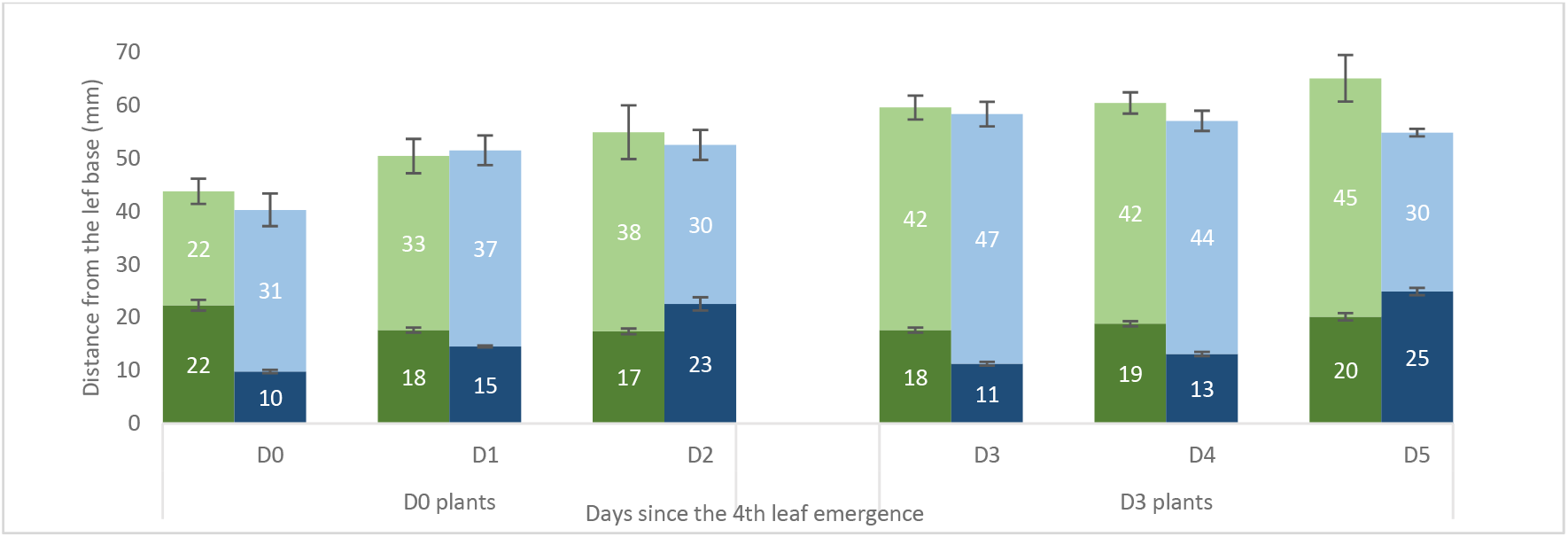
The effect of a 3-day cold period at different developmental stages on the size of meristem, elongation, and growth zone of the maize leaf. The meristem is represented in dark colour and the elongation zone in light colour. The control treatment is represented in green and the cold-treated plants are represented in blue. D0 and D3 correspond to the direct effect of cold, D1 and D4 to the first day of recovery and D2 and D5 to the second day of recovery. Data are presented as mean ± SE, N=6-7. T-test and 2-way Anova results are summarized in **table S1-3**.

Upon cold exposure, the division zone was reduced by 56% (p<0.001) and 36% (p<0.001) for D0 and D3 plants, respectively (**Fig. 7, Table S1**). During the recovery phase meristem size rapidly increased for both D0 and D3 treated plants and after two days the meristem of cold-treated leaves was even larger than that of control plants (30% for D0, p=0.002) and 19% for D3, p=0.011; **Fig. 7, Table S3**).

The size of the growth zone, which includes meristem and elongation zone and is defined by the position where cells reach their mature size, progressively increases after leaf emergence in control conditions (**Fig. 5** and **7**). Exposure to cold has no direct effect on the size of the growth zone but reduces it after 2 days of recovery, particularly in D3-treated plants. This reduction of growth zone size in combination with the increase of the division zone after 2 days of recovery explains the large reduction of the size of the elongation zone (**Fig. 7, Table S3**).

#### Kinematic analysis of cell division and expansion

Kinematic analysis combines measurements of LER (**Fig. 4a),** the cell length profile (**Fig. 6**), and the length of the meristem (**Fig. 7).** Assuming steady-state, cell production rate (P) is normally calculated as the ratio of LER and l_mat_ (Bertels and Beemster, 2020). Due to increasing LER (**Fig. 8a**) and decreasing l_mat_ (**Fig. 8c, Table S1-3**), P progressively increased from D0 to D5 for the control plants (**Fig. 8b; p<0.001)**. Because cell length decreases at the same time (**Fig. 5, 8f)**, cell density in the meristem increases (**Fig. 8h**), this means that cell production needs to be adjusted for the rate of density change over time (Silk, 1992). Adding the rates of density change in the meristem, significantly increases cell production rates, particularly during the first days after emergence (**Fig. 8b**).

**Figure 8:**
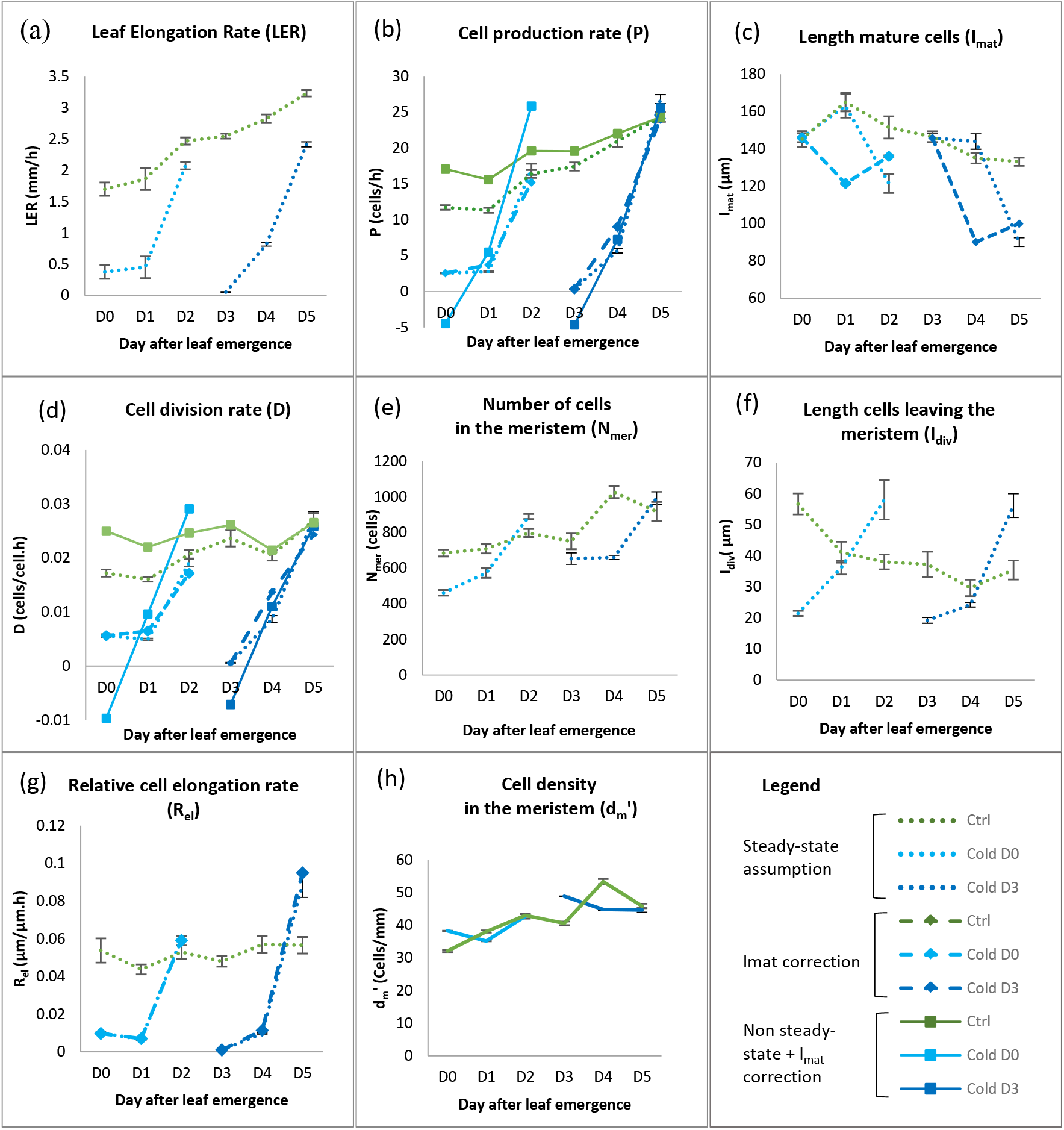
The effect of a 3-day cold period at different developmental stages on cell division and expansion. a. Leaf elongation rate, b. Cell production rate, c. Length of mature cells, d. Cell division rate, e. Number of cells in the meristem, f. Length cells leaving the meristem, g. Relative cell elongation rate, h. Cell density assuming a meristem of 2cm. Plants in control conditions (green line) were harvested from D0 before leaf emergence to D5 after leaf emergence. Cold-treated plants (blue line) were either placed in the cold at D0 before leaf emergence (light blue) or D3 after leaf emergence (dark blue). Those plants were harvested directly after cold (D0 and D3), after 1 day (D1 and D4), and after 2 days of recovery (D2 and D5). Data are represented as mean ± SE (n=6-7). 3-way ANOVA (Cold timing x Harvesting x Treatment) was performed. The data are summarized in **table S4**. In a, e, f and h, values are independent of steady-state assumptions. In panels b, c, d and g, the dotted lines represent calculation assuming steady-state. The dashed lines represent a recalculation of the parameter during the 1^st^ and 2^nd^ days of cold recovery with the corrected l_mat_ from the cell length profile. The full lines correspond to recalculation with corrected l_mat_ and cell density changes in the meristem (**See Method S1 for calculation details**).

Average cell division rates (D) are calculated as cell production rates divided by the number of cells in the meristem. Assuming steady-state, D increased over time. However, using cell production rates corrected for non steady-state showed that D is approximately stable over time. Similar to cell production rates, the largest difference occurred during the earliest stages (**Fig. 8e**). Average cell expansion rates remained approximately stable throughout the development of control plants (**Fig. 8g**).

In D3-exposed plants, the reduction of LER in response to cold was stronger than in plants exposed in D0 (**Fig. 4a & 8a**). Cell division rates based on steady-state cell equations show that this was due to a reduction of cell production rate by 78% and 98% in D0 and D3 exposed plants (**Fig. 8b**). While exposure to cold itself did not affect the cell length profile in the elongation zone (**Fig. 6a,d**), cell elongation was surprisingly reduced by 82% and 98% in plants exposed to cold on D0 and D3, respectively (**Fig. 8g**). The reduced size of the meristem in response to cold (**Fig. 7**) corresponds with a reduction of the number of cells in the meristem by 33% (p<0.001) and 13% (p<0.001) (**Fig. 8e**), despite a minor compensation by shorter cells (**Fig. 6a, d**).

Cell division rates calculated based on steady-state assumptions showed a rapid increase during the first day of recovery for D3 exposed plants, whereas they remain strongly inhibited in D0 exposed plants. After two days of recovery average cell division rates are close to those of control plants for both exposures (**Fig. 8e**). The number of cells in the meristem also progressively increased during recovery and was also fully restored after 2 days of recovery (**Fig. 8e**).

When calculating cell division rates based on (non steady-state) cell production rates (that include density changes in the meristem; full lines in **Fig. 8b**), cell division rates were negative during the cold and higher than the rates based on steady-state assumptions, particularly for D0 treated plants **(Fig. 8d**). This demonstrates the impact of cell density changes in the meristem on the calculations of division rates.

In contrast to cell division rates, cell expansion rates remain inhibited during the first day of recovery, but after 2 days, they were even 11 and 68% higher than in their respective control leaves, suggesting an overcompensation (**Fig. 8g**).

## Discussion

Cold is a key factor affecting the growth and yield of maize in Northern Europe (Ben-Haj-Salah and Tardieu, 1995; Giauffret et al., 1995). Thus, developing more cold-tolerant varieties is crucial for breeders and agronomists. While cold responses have extensively been studied at the physiological, cellular and molecular levels (Burnett and Kromdijk, 2022; Zhou et al., 2022) (Lainé et al., 2022), little is known about the recovery process. Therefore, we characterized the growth response of maize leaf growth during a 3-day cold stress and the first two days of recovery at the whole plant, individual 4^th^ leaf and cellular level.

We demonstrate for the first time a different sensitivity of the same leaf depending on its developmental stage (before or after emergence). We show that during the cold, growth was more severely reduced in emerged (D3 exposed) leaves due to a stronger reduction in cell production and elongation rate than in non-emerged leaves (D0 exposed). Inversely, D3 leaves recovered better (complete recovery of LER) than non-emerged leaves (D0 plants) due to a higher cell division rate after 1 day of recovery (**Fig. 8d**) and a higher cell elongation rate (**Fig. 8g**) after 2 days of recovery.

### Cold differently affects leaves just prior to and after emergence

The leaf elongation rate is first exponential (at leaf initiation) and, once the elongation zone is formed, it becomes linear during a steady-state before declining until reaching its final leaf length (Parent et al., 2009). We hypothesized that cold differently affects the growth of leaves that were in different phases of their development. Consistently, when cold occurs during the declining phase of LER, final leaf length is affected only slightly, i.e., leaf 3 decreased by 5% (**Fig. 2a-b**). Exposure during steady-state growth considerably reduced final leaf length, i.e., leaf 4 is decreased by 14% (**Fig. 2b**). Thereafter, the difference in final leaf length (of the same cold-treated plants) tends to decrease with increasing leaf rank (**Fig. 2b**) (Louarn et al., 2010). Surprisingly, if a leaf is exposed to cold when it was about to emerge, leaf length is most strongly reduced (18%; **Fig. 4b**). This suggests that leaves develop an improved mechanism to recover from cold after their emergence.

We could not observe the effect of cold at the leaf initiation stage, but it has been shown that if cold occurs during the exponential (leaf initiation) phase, the duration of the linear phase is lengthened so that the final size of the leaf is not reduced (Louarn et al., 2010). On the other hand, we observed that if cold occurs during the linear phase of the leaf growth, it is slowed-down during the stress and its final leaf length is reduced (**Fig. 2a-b**). There are two explanations for this. First, we show that similar to observations by Louarn et al. (2010), when cold occurred during its linear phase, the duration of steady-state growth was shortened (arrows in **Fig. 2a**) and the final size was reduced. This is similar to the response of sorghum plants where salinity reduces the leaf growth rate and shortens the period of rapid leaf elongation, thereby producing shorter leaves (Bernstein et al 1993). Secondly, consistent with observations by Birch et al. (1998) and Plancade et al. (2022), leaves from cold-treated plants emerged faster than controls, having a shorter phyllochron. This suggests that the rate of development is increased, which may explain the shorter duration of leaf expansion and, thereby, a shorter final leaf length (**Fig. 3**). This is consistent with the findings of (Riva-Roveda et al., 2016) in maize leaves responding to cold stress and with water stress in rice (Pablo et al., 2022).

### Non Steady-state kinematic growth analysis

In contrast to the frequently assumed steady-state, we observed that in control conditions LER and elongation zone size increase during the first days after leaf emergence, while cell length decreases throughout the growth zone (**Fig. 5**). Contrasting observations were made in rice, where the size of the elongation zone decreases and LER starts declining after leaf emergence (Parent et al., 2009). When steady-state growth is not met, parameters derived from the kinematic method (Sprangers et al., 2016) must be revised. Our results show that increasing leaf elongation rate in control conditions is driven by increased cell production in the meristem, which is partly offset by decreasing mature cell length. An important observation is the decreasing size of proliferating cells, increasing the density and number of dividing cells. This led us to calculate cell production and cell division rates by adding the rate of density change in the meristem (Silk, 1992). Our results show that during the first days of emergence, this has a substantial effect, but at 4-5 days after emergence, when we usually perform kinematic analyses based on steady-state assumptions, the difference becomes negligible (**Fig. 8b,d).**

Non steady-state responses, however, are more pronounced during recovery from cold. This is immediately obvious from the dynamic changes in the cell length profile, which show a differential effect of cold on the subsequent cell expansion of cells in different stages of their development. Rapid changes in cell density in the meristem also considerably affected cell production and division rates (Fig. 8b and d). Moreover, time-related parameters such as cell cycle duration, time cells spend in the division and elongation that are reliably estimated under steady-state conditions (Sprangers et al., 2016; Bertels and Beemster, 2020), become irrelevant in the context of dynamically changing conditions.

### Active suppression of growth controlled by cell division and expansion

After 3 days of cold, LER decreased by 98% by 78% for D3 and D0 plants, respectively (**Table S1**) due to a reduction of cell production and cell elongation (**Fig. 8b, g**). Similar effects on cell production rate were also observed in other abiotic stresses such as salt stress (West et al., 2004), drought stress (Avramova et al., 2015) and light and temperature stress (Granier and Tardieu, 2000). This was also observed when cold is limited to the night period (25°C day/ 4°C night cycles), where cold only affected the activity of the basal meristem of the leaves (Rymen et al., 2007). Transcriptome analyses showed that the growth response to cold stress is mediated, amongst other processes, by altered expression of cell cycle genes (Rymen et al., 2007). In contrast to our findings, when cold only occurs during the night, the number of dividing cells in the meristem increased in cold-treated leaves in comparison to the control (Rymen et al., 2007). However, the shortening of cell length in the division zone in response to our treatment closely correlates with similar observations by Rymen et al 2007.

Cold also led to a reduced mature cell length for both D3 and D0 treated plants. Due to the time it takes for cells to move through the growth zone, this effect could only be observed from the cell length profile during the recovery period (**Fig. 6**). Mature cell length is often not affected by direct abiotic stresses in maize leaves as a lower cell expansion rate is often compensated by a longer time cell spends in the elongation zone (Rymen et al., 2007; Avramova et al., 2015; Fina et al., 2017).To our knowledge, this study demonstrates for the first time a transient reduction in mature cell length upon recovery (**Fig. 6)**. Because under these conditions cell length profile is not in steady-state, the effect of cold stress on cell expansion only becomes indirectly visible after two days, when expanding cells reach maturity. One plausible reason for the disruption of cell expansion is a modification of the cell wall into a more rigid structure leading to a restriction in cell growth by limitation of cell expansion (Podgórska et al., 2017). Accordingly, Riva-Roveda et al (2016) reported that a reduction in cell elongation was responsible for 8 times the decrease in leaf growth during cold stress (10°C day/7°C night). In addition, the decrease in cell elongation rate was underpinned by the downregulation of gene coding for expansin protein (Riva-Roveda et al., 2016). Expansins are known to mediate cell wall loosening by non-enzymatically triggering a pH-dependent relaxation which enables cell expansion (Marowa et al., 2016). Our results provide a basis to determine in much more detail at which stage of cellular development, i.e. where in the growth zone, such changes are occurring.

### Recovery is more important than direct response to cold

Our results show that LER, cell production and cell expansion rates are more affected by cold once the leaf has emerged. D0 plants were less affected during the chilling stress with LER decreasing by 78% than D3 plants where LER decreased by 98% (**Table S1**). A plausible reason for that is that non-emerged leaves are still surrounded by the whorl of older leaves and consequently are better protected. Surprisingly, D3 plants recover better, resulting in a smaller effect on final leaf length (**Fig. 4b**). An explanation for this is that plants enter in a standby mode during cold stress as a strategy to recover better. This suggestion was also underlined by Riva-Roveda et al., (2016) in maize leaf with cold stress and Clauw et al (2015) in Arabidopsis leaf with severe drought stress. Following this reasoning, it would suggest that D3 plants can recover better than D0 due to a better growth cessation during cold. Indeed, increasing evidence indicates that plants actively repress growth under stress conditions as an adaptive strategy to maximize recovery (Chapin, 1991; Kasuga et al., 1999).

After cold stress, an innate recovery response is activated, losing primary stress symptoms and providing plant regeneration through further growth and development of plants (Hasanfard et al., 2021). Our results revealed that the most efficient recovery occurred in the already emerged leaf. A recent review on metal stress in plants suggests that recovering faster would be more efficient than a slow one (Chmielowska-Bąk and Deckert, 2021). It is important to notice that, in our study, D3 plants were capable of full growth recovery after three days in control conditions (**Fig. 4a**). In contrast, even after five days of growth in control conditions, D0 treated leaves reached a maximum of 78% of LER of the corresponding controls (**Fig. 4a**). At the cellular level, the fast and complete recovery observed in D3 plants was due to higher cell division on the 1^st^ day of recovery and higher cell elongation rate after 2-days of recovery (**Fig. 8d, g**), but regarding underlying molecular and physiological mechanisms, we can only speculate. One observation is that when exposed to sudden cold stress, maize seedlings exhibit symptoms of drought stress due to an imbalance between transpiration and water uptake (Aroca et al., 2003). Munns et al (2000) demonstrated that the rapid recovery of elongation rate on relief of salt stress was due to the increase in water status. Similarly, Ben Haj Salah and Tardieu (1996) show that an increase in vapour pressure causing a decrease in leaf water potential of about 700 kPa caused a decline and then partial recovery of leaf elongation rate in maize plants. This suggests that the observed recovery difference between emerged and non-emerged leaves may be related to water relations, which result in differences in turgor driving cell expansion.

## Conclusion

Our results demonstrate that final leaf length of non-emerged leaves (D0) was more affected by cold than the emerged ones (D3). This was not the result of better growth of D3 leaves during cold, but a faster and complete recovery of their leaf elongation rate. At the cellular level, this difference was due to a stronger inhibition of division and expansion during the stress, and subsequently a higher cell division rate on the 1st and a higher cell elongation rate on the 2nd-day recovery respectively. The recovery period is more determinant for the impact on final leaf size than the tolerance during cold stress.

## Acknowledgements

This work has been supported by The University Research Fund (BOF) with a PhD fellowship We thank Senne Note who contributed to data generation.

## Author Contributions

CL performed the data analysis and wrote the paper.

HAE and GB conceived the idea, supervised the analyses, and contributed to the writing.

## Supporting Information

Additional Supporting Information may be found online in the Supporting Information section at the end of the article.

| Filename | Description |
| --- | --- |
| **Figure S1-3,**<br>PDF document,<br>119 KB | **Fig. S1** Time course in days of leaf length (1-9) maize plants in control or cold condition for 3 days at D3 after 4<sup>th</sup> leaf emergence.<br>**Fig. S2** Time course in days of leaf elongation rate from the 4<sup>th</sup> leaf of maize when grown fully in control condition or placed for 3 days in cold at D0 before 4<sup>th</sup> leaf emergence or at D3 after 4<sup>th</sup> emergence and transferred back to control condition for recovery<br>**Fig. S3** The effect of a 3-day cold spell on the cell length profile of epidermal cells in the growth zone of the maize leaf after 3 days of recovery. |
| **Method S1,**<br>PDF document,<br>318 KB | **Method S1** Explanation for correction of the kinematic parameters when leaf growth is in non steady-state |
| **Table S1-4,**<br>PDF document,<br>150 KB | **Table S1** A 2-way ANOVA analysis of kinematic parameters comparing the effect of cold with control (D0 vs D3).<br>**Table S2** A 2-way ANOVA analysis of kinematic parameters comparing the effect of cold after 1 day recovery with control (D1 vs D4).<br>**Table S3** A 2-way ANOVA analysis of kinematic parameters comparing the effect of cold after 2 days recovery with control (D2 vs D5).<br>**Table S4** A 3-way ANOVA analysis of kinematic parameters (treatment x harvesting time x cold timing) |

## References

Ade-Ademilua OE, Botha CEJ, Strasser RJ (2005) A re-evaluation of plastochron index determination in peas — a case for using leaflet length. South Afr J Bot 71: 76–80

Allen DJ, Ort DR (2001) Impacts of chilling temperatures on photosynthesis in warm-climate plants. Trends Plant Sci 6: 36–42

Aroca R, Vernieri P, Irigoyen JJ, Sánchez-Dıéz M, Tognoni F, Pardossi A (2003) Involvement of abscisic acid in leaf and root of maize (Zea mays L.) in avoiding chilling-induced water stress. Plant Sci 165: 671–679

Atkinson LJ, Sherlock DJ, Atkin OK (2015) Source of nitrogen associated with recovery of relative growth rate in Arabidopsis thaliana acclimated to sustained cold treatment. Plant Cell Environ 38: 1023–1034

Avila LM, Obeidat W, Earl H, Niu X, Hargreaves W, Lukens L (2018) Shared and genetically distinct Zea mays transcriptome responses to ongoing and past low temperature exposure. BMC Genomics 19: 761

Avramova V, AbdElgawad H, Zhang Z, Fotschki B, Casadevall R, Vergauwen L, Knapen D, Taleisnik E, Guisez Y, Asard H, et al (2015) Drought Induces Distinct Growth Response, Protection, and Recovery Mechanisms in the Maize Leaf Growth Zone. Plant Physiol 169: 1382–1396

Aydinoglu F (2020) Elucidating the regulatory roles of microRNAs in maize (Zea mays L.) leaf growth response to chilling stress. Planta 251: 38

Beemster GTS, Baskin TI (1998) Analysis of Cell Division and Elongation Underlying the Developmental Acceleration of Root Growth in Arabidopsis thaliana. Plant Physiol 116: 1515–1526

Beemster GTS, Masle J (1996) The role of apical development around the time of leaf initiation in determining leaf width at maturity in wheat seedlings (Triticum aestivum L.) with impeded roots. J Exp Bot 47: 1679–1688

Ben Haj Salah H, Tardieu F (1996) Quantitative analysis of the combined effects of temperature, evaporative demand and light on leaf elongation rate in well-watered field and laboratory-grown maize plants. J Exp Bot 47: 1689–1698

Ben-Haj-Salah H, Tardieu F (1995) Temperature Affects Expansion Rate of Maize Leaves without Change in Spatial Distribution of Cell Length (Analysis of the Coordination between Cell Division and Cell Expansion). Plant Physiol 109: 861–870

Bernstein N, Lauchli A, Silk WK (1993) Kinematics and Dynamics of Sorghum (Sorghum bicolor L.) Leaf Development at Various Na/Ca Salinities (I. Elongation Growth). Plant Physiol 103: 1107–1114

Bertels J, Beemster GTS (2020) leafkin—An R package for automated kinematic data analysis of monocot leaves. Quant Plant Biol. doi: 10.1017/qpb.2020.3

Birch C, Vos J, Kiniry J, Bos HJ, Elings A (1998) Phyllochron responds to acclimation to temperature and irradiance in maize. Field Crops Res 59: 187–200

Brüggemann W, van der Kooij TAW, van Hasselt PR (1992) Long-term chilling of young tomato plants under low light and subsequent recovery: I. Growth, development and photosynthesis. Planta 186: 172–178

Burnett AC, Kromdijk J (2022) Can we improve the chilling tolerance of maize photosynthesis through breeding? J Exp Bot erac045

Chapin FS III (1991) Integrated Responses of Plants to Stress: A centralized system of physiological responses. BioScience 41: 29–36

Chmielowska-Bąk J, Deckert J (2021) Plant Recovery after Metal Stress—A Review. Plants 10: 450

Cholakova R, Vassilev A (2017) Effect Of Chilling Stress On The Photosynthetic Performance Of Young Plants From Two Maize (Zea Mays) Hybrids. CBU Int Conf Proc 5: 1118–1123

Clauw P, Coppens F, De Beuf K, Dhondt S, Van Daele T, Maleux K, Storme V, Clement L, Gonzalez N, Inzé D (2015) Leaf Responses to Mild Drought Stress in Natural Variants of Arabidopsis. Plant Physiol 167: 800–816

De Vos D, Nelissen H, AbdElgawad H, Prinsen E, Broeckhove J, Inzé D, Beemster GTS (2020) How grass keeps growing: an integrated analysis of hormonal crosstalk in the maize leaf growth zone. New Phytol 225: 2513–2525

DeRidder BP, Crafts-Brandner SJ (2008) Chilling stress response of postemergent cotton seedlings. Physiol Plant 134: 430–439

Erickson RO (1976) Modeling of Plant Growth. Annu Rev Plant Physiol 27: 407–434

Farooq M, Aziz T, Wahid A, Lee D-J, Siddique KHM (2009) Chilling tolerance in maize: agronomic and physiological approaches. Crop Pasture Sci 60: 501–516

Fina J, Casadevall R, AbdElgawad H, Prinsen E, Markakis MN, Beemster GTS, Casati P (2017) UV-B Inhibits Leaf Growth through Changes in Growth Regulating Factors and Gibberellin Levels. Plant Physiol 174: 1110–1126

Fiorani F, Beemster G (2006) Quantitative analyses of cell division in plants. PLANT Mol Biol 60: 963–979

Frei O (2000) Changes in yield physiology of corn as a result of breeding in northern Europe. Maydica 45: 173–183

Garbero M, Andrade A, Reinoso H, Fernández B, Cuesta C, Granda V, Escudero C, Abdala G, Pedranzani H (2012) Differential effect of short-term cold stress on growth, anatomy, and hormone levels in cold-sensitive versus -resistant cultivars of Digitaria eriantha. Acta Physiol Plant 34: 2079–2091

Giauffret C, Bonhomme R, Derieux M (1995) Genotypic differences for temperature response of leaf appearance rate and leaf elongation rate in field-grown maize. Agron. 2 15 123–137 1995

Gómez LD, Vanacker H, Buchner P, Noctor G, Foyer CH (2004) Intercellular Distribution of Glutathione Synthesis in Maize Leaves and Its Response to Short-Term Chilling. Plant Physiol 134: 1662–1671

Granier C, Tardieu F (1998) Is thermal time adequate for expressing the effects of temperature on sunflower leaf development? Plant Cell Environ 21: 695–703

Granier C, Tardieu F (2000) Sunflower leaf growth under changing environmental conditions. OCL - Ol Corps Gras Lipides 7: 219–228

Greaves JA (1996) Improving suboptimal temperature tolerance in maize- the search for variation. J Exp Bot 47: 307–323

Green PB (1976) Growth and Cell Pattern Formation on an Axis: Critique of Concepts, Terminology, and Modes of Study. Bot Gaz 137: 187–202

Hasanfard A, Rastgoo M, Izadi Darbandi E, Nezami A, Chauhan BS (2021) Regeneration capacity after exposure to freezing in wild oat (Avena ludoviciana Durieu.) and turnipweed (Rapistrum rugosum (L.) All.) in comparison with winter wheat. Environ Exp Bot 181: 104271

Kasuga M, Liu Q, Miura S, Yamaguchi-Shinozaki K, Shinozaki K (1999) Improving plant drought, salt, and freezing tolerance by gene transfer of a single stress-inducible transcription factor. Nat Biotechnol 17: 287–291

Lainé CMS, AbdElgawad H, Beemster GTS (2023) A meta-analysis reveals differential sensitivity of cold stress responses in the maize leaf. Plant Cell Environ. doi: 10.1111/pce.14608

Louarn G, Andrieu B, Giauffret C (2010) A size-mediated effect can compensate for transient chilling stress affecting maize (Zea mays) leaf extension. New Phytol 187: 106–118

Marowa P, Ding A, Kong Y (2016) Expansins: roles in plant growth and potential applications in crop improvement. Plant Cell Rep 35: 949–965

Meicenheimer RD (2014) The plastochron index: still useful after nearly six decades. Am J Bot 101: 1821–1835

Miedema P (1982) The Effects of Low Temperature on Zea mays. In NC Brady, ed, Adv. Agron. Academic Press, pp 93–128

Muller B, Reymond M, Tardieu F (2001) The elongation rate at the base of a maize leaf shows an invariant pattern during both the steady-state elongation and the establishment of the elongation zone. J Exp Bot 52: 1259–1268

Munns R, Passioura JB, Guo J, Chazen O, Cramer GR (2000) Water relations and leaf expansion: importance of time scale. J Exp Bot 51: 1495–1504

Pablo J, Clerget B, Bueno C, Dionora J, Domingo A, Guzman C, Aguilar E, Cadiz N, Sta. Cruz P (2022) Phyllochron duration and changes through rice development shape the vertical leaf size profile. doi: 10.1101/2022.03.12.484079

Parent B, Conejero G, Tardieu F (2009) Spatial and temporal analysis of non-steady elongation of rice leaves. Plant Cell Environ 32: 1561–1572

Pettigrew W (2002) Improved Yield Potential with an Early Planting Cotton Production System. Agron J. doi: 10.2134/agronj2002.0997

Plancade S, Marchadier E, Huet S, Ressayre A, Nous C, Dillmann C (2022) A new hypothesis-testing model for phyllochron based on a stochastic process - application to analysis of genetic and environment effects in maize. bioRxiv

Podgórska A, Burian M, Gieczewska K, Ostaszewska-Bugajska M, Zebrowski J, Solecka D, Szal B (2017) Altered Cell Wall Plasticity Can Restrict Plant Growth under Ammonium Nutrition. Front. Plant Sci. 8:

Rankenberg T, Geldhof B, Veen H van, Holsteens K, Poel BV de, Sasidharan R (2021) Age-Dependent Abiotic Stress Resilience in Plants. Trends Plant Sci 26: 692–705

Riva-Roveda L, Escale B, Giauffret C, Périlleux C (2016) Maize plants can enter a standby mode to cope with chilling stress. BMC Plant Biol 16: 212

Rymen B, Fiorani F, Kartal F, Vandepoele K, Inzé D, Beemster GTS (2007) Cold Nights Impair Leaf Growth and Cell Cycle Progression in Maize through Transcriptional Changes of Cell Cycle Genes. Plant Physiol 143: 1429–1438

Salesse-Smith CE, Sharwood RE, Busch FA, Stern DB (2020) Increased Rubisco content in maize mitigates chilling stress and speeds recovery. Plant Biotechnol J 18: 1409– 1420

Silk WK (1992) Steady Form from Changing Cells. Int J Plant Sci 153: S49–S58

Sprangers K, Avramova V, Beemster GTS (2016) Kinematic Analysis of Cell Division and Expansion: Quantifying the Cellular Basis of Growth and Sampling Developmental Zones in Zea mays Leaves. JoVE J Vis Exp e54887

Toda K, Takahashi R, Iwashina T, Hajika M (2011) Difference in chilling-induced flavonoid profiles, antioxidant activity and chilling tolerance between soybean near-isogenic lines for the pubescence color gene. J Plant Res 124: 173–182

Vilonen L, Ross M, Smith MD (2022) What happens after drought ends: synthesizing terms and definitions. New Phytol 235: 420–431

Wang JY (1960) A Critique of the Heat Unit Approach to Plant Response Studies. Ecology 41: 785–790

West G, Inzé D, Beemster GTS (2004) Cell cycle modulation in the response of the primary root of Arabidopsis to salt stress. Plant Physiol 135: 1050–1058

Yeung E, van Veen H, Vashisht D, Sobral Paiva AL, Hummel M, Rankenberg T, Steffens B, Steffen-Heins A, Sauter M, de Vries M, et al (2018) A stress recovery signaling network for enhanced flooding tolerance in Arabidopsis thaliana. Proc Natl Acad Sci 115: E6085–E6094

Zhou X, Muhammad I, Lan H, Xia C (2022) Recent Advances in the Analysis of Cold Tolerance in Maize. Front Plant Sci 13: 866034

